# Transpiration, Photoinhibition and Non-photochemical Quenching Reciprocally Control Foliar Heat Emission

**DOI:** 10.1101/2025.05.27.656286

**Authors:** Roshanak Zarrin Ghalami, Maria Duszyn, Muhammad Kamran, Paweł Burdiak, Piotr Gawroński, Stanisław M. Karpiński

## Abstract

Global warming intensifies heat waves and drought, causing energy absorption in excess (EAE) in plants, thereby inhibiting transpiration and photosynthesis. Non-photochemical quenching (NPQ) contributes significantly to EAE dissipation as heat, with leaves emitting ca. 107 W m^-2^ while absorbing ca. 417 W m^-2^. Foliar temperature is further elevated when transpiration is inhibited, but NPQ is not increased. Surprisingly, during generic photosynthetic electron transport and NPQ partial inhibition, foliar temperature under EAE was reduced considerably. Our findings emphasize the importance of mutual co-regulation of photosynthetic electron transport, NPQ, and transpiration in foliar temperature regulation and suggest that overall heat emission from forest ecosystems may conditionally vary on a terawatt (TW) scale per 1,000,000 km^-2^. Global warming models do not take into account this putative phenomenon.

**One Sentence Summary:** Photosynthesis and Foliar Thermoregulation

## Main Text

Terrestrial plants are estimated to contribute up to 20% of global photosynthesis, thereby playing a vital role in mitigating global warming by photosynthetic CO_2_ assimilation and simultaneously cooling the surrounding air through transpiration and evaporation (*1, 2*). The year 2024 was recorded as the warmest period since the onset of global warming, with the rainforests in the Amazon, central Africa, and Southeast Asia experiencing severe drought stress, which, in consequence, caused record-high wildfires. According to the data from the Fire Program of the National Institute for Space Research (INPE), 137,538 fire outbreaks in the Amazon were recorded by the beginning of December 2024, an increase of 43% compared to 2023. Fires in the forest ecosystems have led to a catastrophic deterioration in air quality and further acceleration of the greenhouse effect. Furthermore, global warming triggers extreme weather events in other regions that increase both the frequency and intensity of heavy rainfall (*3*), leading to flooding, which plants experience as hypoxia. These environmental factors reduce both transpiration and photosynthesis (*4*), raising concerns that terrestrial photosynthesis and CO_2_ assimilation could collapse under the increasing greenhouse effect. Therefore, understanding the interplay between global warming, transpiration, non-photochemical quenching (NPQ), and photosynthesis in plant ecosystems is critical, especially as deforestation and desertification continue to accelerate. Modeling studies predict that desert areas could expand to cover up to 10% of the Earth’s land surface by 2050 (*5, 6*), potentially leading to further declines in global photosynthesis (*7*).

In essence, light energy absorbed by photosystems is separated into three main energy channels: fluorescence, heat and photochemistry (*8*). Under non-saturating light conditions, light-harvesting complexes demonstrate exceptional efficiency, with up to 99% of absorbed photons utilized in electrical charge separation (*8–11*). However, when plants experience EAE, a photosynthetic P680 reaction center quickly saturates. This reduces the amount of energy allocated to photochemistry, increasing the energy dissipated as heat (*12–16*).NPQ, a key mechanism for energy dissipation as heat (qE), is typically assessed by chlorophyll fluorescence measurements. Therefore, NPQ does not directly quantify foliar heat emission and changes in leaf temperature (*8–16*). This is a significant knowledge gap because the extent of heat radiated from leaves during EAE remains largely unknown, with only limited estimation and theoretical predictions available (*12–16*). Therefore, it is commonly believed that plants cool down their surroundings through transpiration and CO_2_ assimilation (*17*), thus reducing the progression of global warming. However, in suboptimal conditions, e.g., high solar irradiation, high temperature and transient drought or flooding, transpiration and CO_2_ assimilation are inhibited, thus increasing the need for EAE dissipation as heat.

In sunlight plants naturally absorb more energy than the amount required for photochemistry (*8, 9*). Therefore, they employ diverse strategies to survive and maintain optimal photosynthesis under energy absorption in excess (EAE) usually combined with drought and high-temperature stress. The mechanisms regulating the fate of absorbed EAE include NPQ, in which the energy-dependent quenching (qE), zeaxanthin-dependent quenching (qZ), carotenoids, and singlet oxygen-dependent quenching are major compounds (*14–16, 18–20*). However, increased wave-like, discrete and spatial NPQ changes in directly stressed leaves as well as in leaves undergoing systemic and network acquired acclimation (SAA and NAA) are also observed due to electrical and reactive oxygen species (ROS) wave-like signaling (*21–24*). This indicates that NPQ is not only involved in EAE dissipation as heat but also is involved in SAA and NAA retrograde signaling mechanisms. If plants cannot cope with EAE they finally induce cell death as the ultimate end of photosynthesis, respiration, and cell cycle (*22–27*).

We previously demonstrated that leaves of field-grown transgenic hybrid Aspen trees with deregulated (reduced) MPK4 protein levels showed higher NPQ and foliar temperature, which was accompanied by increased ROS content and resulted in enhanced foliar cell death and reduced growth (*13*). The other reports clearly demonstrated that specific transgenic amelioration of NPQ and violaxanthin–antheraxanthin–zeaxanthin (VAZ) cycle enzymes significantly improved yield in field-grown crop plants due to faster recovery of photosynthesis after transition from sunlight to shadow (*18, 28*). In another article, we revealed that physiological light memory, ROS content, and foliar cell death in response to EAE and UV directly depend on NPQ level (*24*). It is also known that photorespiration mediates chloroplast retrograde signaling for cell death, and specific amelioration of the photorespiratory pathway leads to increased crop yield (*25, 29*).

Our studies used a novel methodology for measurements of mutual mechanisms of foliar thermoregulation, which revealed that plant leaves may conditionally cool down or warm up their surroundings.

## Results

### Impact of EAE on foliar temperature - spatial and developmental patterns

To get a clue about spatial and developmental patterns of foliar thermoregulation during EAE in model plant species, we directly measured Arabidopsis leaves temperature using a thermal camera, as described previously (*12, 13*). In the middle of the photoperiod, the 5-week-old plants were exposed to EAE with photon flux density (PFD) of 2000 ±50 μmol m^-2^ s^-1^ for 180 s, and subsequent changes in foliar temperature were recorded (Fig. 1A-C, Video S1 and S2). Following excess light exposure, significant increases in foliar temperature (ΔT, defined as the change relative to the initial baseline in ambient light) were observed in both juvenile and mature leaves, but with a more pronounced rise in mature leaves. Interestingly, foliar temperature dynamics during EAE is complex and after 180 seconds illumination, the maximum temperature was reached, with a ΔT of 13.1 ±0.7 °C in the apical region and 16.5 ±0.5 °C in the mature leaves (Fig. 1B). Upon cessation of the excess light, leaf temperature rapidly decreased, however foliar temperature after 30 s in ambient light was ca. 5 °C higher than at the start of the experiment. Interestingly, foliar temperature regulation within one leaf is discrete, emergent, dynamic, and spatial (wavy-like) (Fig. 1C, Video S1 and S2).

**Figure 1.**
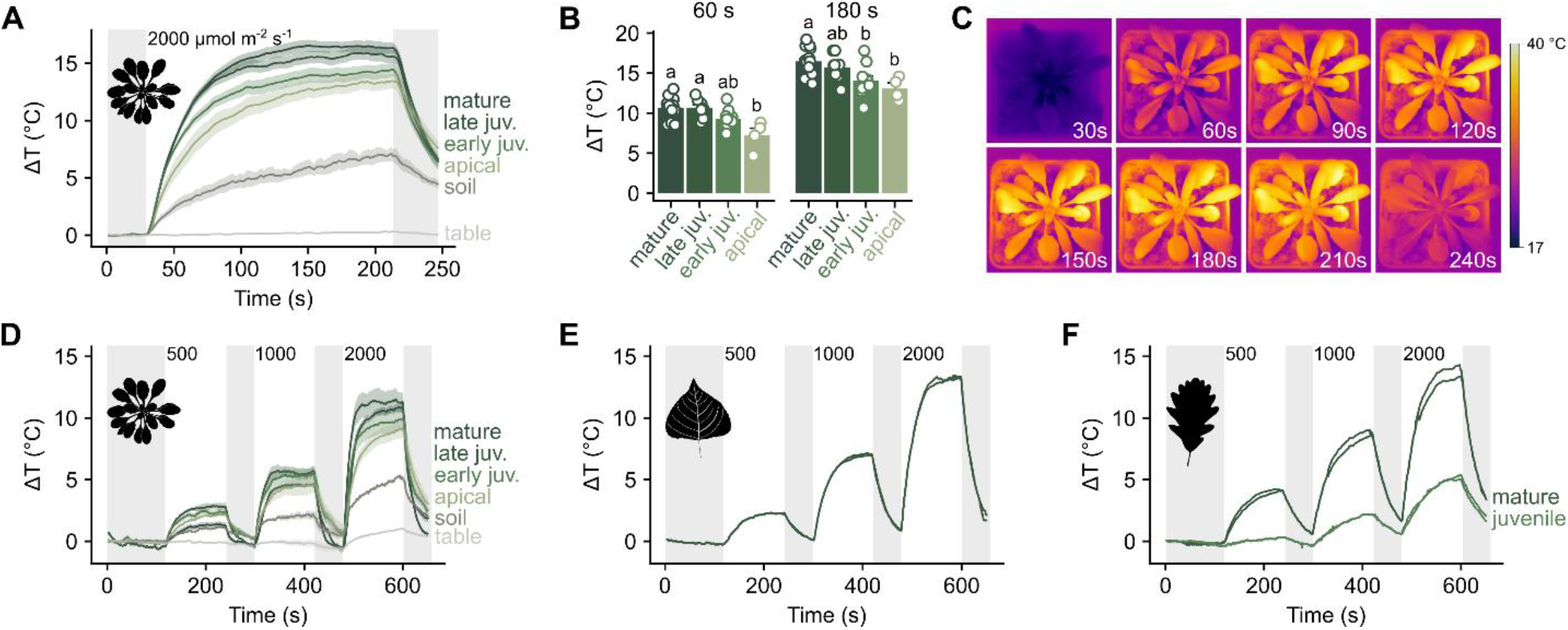
Light intensity-dependent variations in foliar temperature across different plant species and leaf developmental stages. (**A**) Temporal profile of foliar temperature change (ΔT) in *Arabidopsis thaliana* leaves exposed to photon flux density (PFD) of 2000 μmol m^−2^ s^−1^. Leaves were acclimated to laboratory room temperature under PFD ca. 50 μmol m^−2^ s^−1^ LED light. After 30 seconds of initial foliar temperature measurements, leaves were exposed to PFD of 2000 μmol m^−2^ s^−1^ LED light for 180 seconds. Data represent the mean ± SEM from four independent experiments (measuring in total up to 13 different leaf areas). (**B**) Selected time points extracted from the ΔT profile (panel A) in different developmental stages of Arabidopsis leaves. Letters over the bars indicate homogeneous groups based on a Tukey HSD test (p < 0.05). (**C**) Representative thermographic images of an Arabidopsis rosette, illustrating dynamic temperature changes. (**D–F**) Foliar temperature responses (ΔT) of Arabidopsis (**D**), hybrid aspen (*Populus tremula × P. tremuloides*) (**E**), and *Quercus* sp. (**F**) subjected to increasing PFD (500, 1000, and 2000 μmol m^−2^ s^−1^). For these panels, each light intensity was applied for 120 seconds, interleaved with 60 seconds of PFD ca. 50 μmol m^−2^ s^−1^, following an initial temperature measurement for a 120 second period. In panel **D**, lines represent mean ±SEM from two independent experiments, whereas panels **E** and **F** display data from several individual experiments.

Subsequently, to get a clue if an increase in foliar temperature corelated with increase in light intensity we exposed the plants to increasing PFD (500, 1000, and 2000 μmol m^−2^ s^−1^) for 120 s at each intensity, with 60 s actinic light intervales between various EAE exposures (Fig. 1D). The ΔT increased with light intensity almost linearly, with the lowest values at 500 μmol m^−2^ s^−1^ (ΔT of 3.1 ±0.2 °C), higher at 1000 μmol m^−2^ s^−1^ (ΔT of 5.8 ±0.3 °C) and the highest at 2000 μmol m^−2^ s^−1^ (ΔT of 13.1 ±0.4 °C). The Arabidopsis results were clear in laboratory conditions, which are different from the outdoor natural conditions in terms of visible and invisible (UV, IR) light quality, intensity and photoperiod, temperature, humidity changes, and various biotic and abiotic stimuli. Therefore, we supposed that foliar thermoregulation in response to an increase in EAE could be different (for example, less dynamic) in leaves growing outdoors. Surprisingly, we also observed that ΔT varied with the leaf age in a manner similar to that described above (i.e. the younger leaves the lower ΔT) and a similar or even more dynamic response to increasing EAE occurs in leaves of *Quercus* sp. and hybrid Aspen (*Populus tremula x P. tremuloides*) using the same protocol (Fig. 1E, F). The foliar temperature responses of both *Quercus* sp. and hybrid aspen growing in the nature resembled those of Arabidopsis in the laboratory (growing chambers). Notably, in *Quercus* sp. ΔT varied with developmental stage, and the difference in ΔT between mature and juvenile leaves was even more pronounced than in Arabidopsis (Fig. 1D, F). Collectively, these results indicate that increasing EAE induces a significant and rapid (dynamic) increase in foliar temperature upon EAE exposure, with mature leaves exhibiting a stronger thermal response compared to juvenile leaves. Moreover, foliar temperature changes within one leaf are discrete and spatial–wave–like (Videos S1 and S2).

### Generic inhibition of transpiration increases foliar temperature

It is generally accepted that terrestrial plants cool down the surrounding environment through transpiration (*17*). However, what happens when transpiration is inhibited and EAE increases is not well understood. To get a closer insight into this process, we measured the temperature of *Acer platanoides* (Maple) leaves during photosynthesis, both under normal natural conditions and when transpiration was artificially inhibited to mimic stomatal closure, but without detaching the leaves from the tree. In addition, we compared leaves from sunny and shaded sides of the tree in June and October 2024 (Fig. 2A, B). On the sunny side, foliar temperatures were 30.0 ± 1.1 °C in June and 26.1 ± 0.7 °C in October, whereas on the shaded side, the temperatures were 22.7 ± 1.3 °C and 20.5 ± 0.8 °C, respectively—supporting our previous observation that foliar temperature increases with light intensity. Air temperature during measurements was 26 °C in June and 20.5 °C in October.

**Figure 2.**
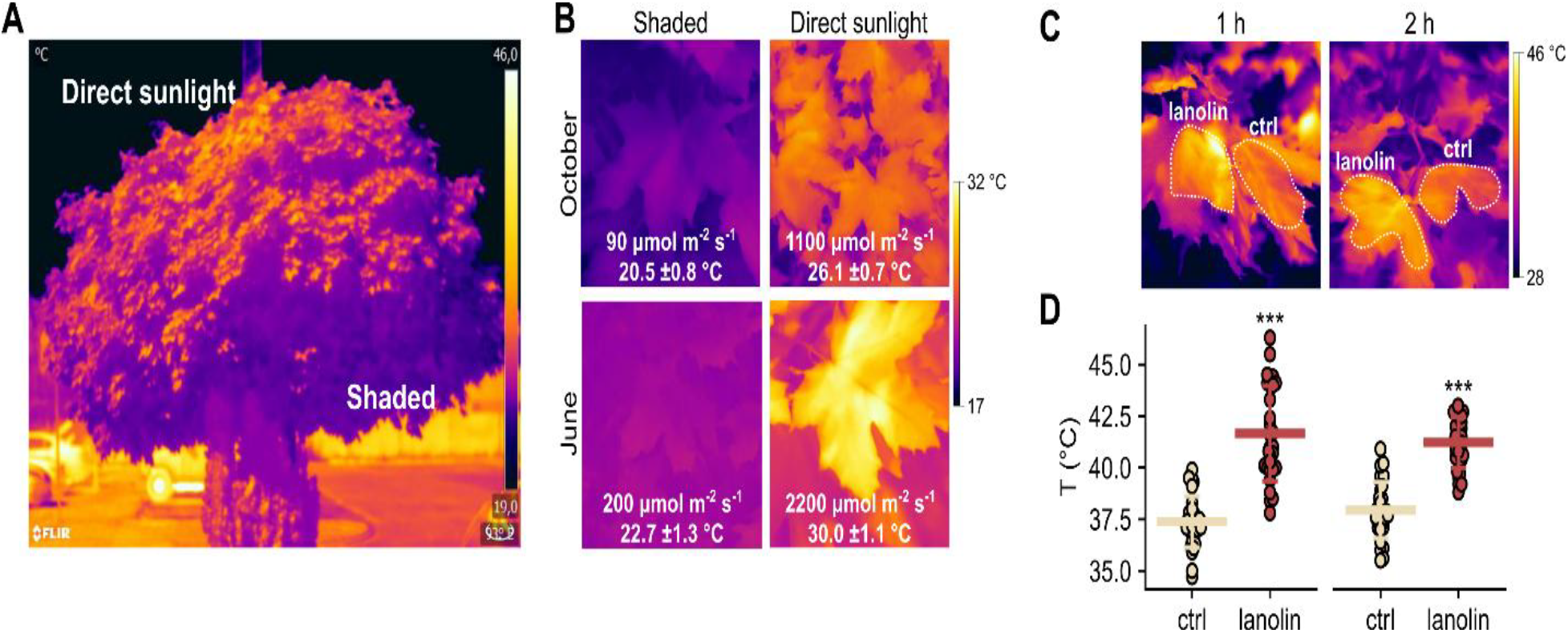
Temperature of sunlight-exposed and shadowed *Acer platanoides* tree leaves in field conditions with not inhibited and with artificially inhibited transpiration. (**A**) Thermographic images of an *Acer platanoides* tree, highlighting the temperature differences between sunny and shaded regions. (**B**) Leaf temperature measurements from shaded and sunny areas, taken at different time points of seasons (June and October). (**C**) Thermographic images and (**D**) corresponding temperature data for control (ctrl) and lanolin-coated not detached leaves after 1 and 2 hours of sunlight exposure. Data represent the mean ±SD. Statistical significance was determined using a t-test (*** *P* < 0.001). For each variant, three leaves were analyzed with 10 point measurements *per* leaf (*n*=30) from three different areas of the tree.

To determine how stomatal conductance influences foliar temperature in non-detached tree leaves, we conducted an experiment using lanolin to artificially restrict transpiration. The abaxial surfaces of selected maple leaves were coated with a smear of lanolin to impede the stomatal conductance and transpiration (*25*), while neighboring leaves with intact stomatal function served as controls. Over a two-hour period of transpiration inhibition, the temperature of lanolin-coated leaves increased by 3.3 – 4.3 °C compared to controls (Fig. 2C, D). At the same time, NPQ and *Fv/Fm* did not significantly change after 1 h but were reduced after 2 h (Table. S1). Foliar average temperature in lanolin-treated leaves was slightly reduced after 2 h in comparison to 1 h after treatment, but was significantly higher than in control leaves (Fig. 2A and B, Table S1). These results indicate that stomatal closure and the subsequent transient inhibition of transpiration, with a significant but small reduction in *F*_v_/*F*_m_ and NPQ, contribute to increased foliar temperature.

### Foliar heat emission from leaves during EAE

To directly quantify EAE dissipated from leaves as heat, we developed and employed a custom-built photo-calorimeter (Fig. S1). This device features a thermally insulated, two-chamber system filled with distilled water, equipped with highly sensitive temperature sensors and an integrated light source for controlled illumination. Temperature differences between the control chamber (without leaves) and the measurement chamber (containing leaves) are continuously recorded. This design allows for the precise detection of subtle water temperature changes induced by heat emitted from the leaf. Due to continuous monitoring of temperature changes over time in both chambers, the photo-calorimeter provides direct, real-time quantification of leaf heat emission under fixed light PAR. Importantly, these measurements were performed under conditions of restricted transpiration, as the leaves were fully submerged in distilled water. This approach complements chlorophyll fluorescence measurements by offering an independent assessment of thermal energy release, which is particularly useful for assessing the contribution of NPQ to overall foliar heat production.

We measured the heat emitted by leaves of laboratory-grown (growth chamber) *Arabidopsis thaliana* and *Euphorbia pulcherrima*, as well as leaves collected from both sunny and shaded sites of *Acer platanoides* tree grown under natural conditions (Table 1). The results indicate that all samples generated comparable amounts of heat (0.149–0.167 cal s^-1^ g^-1^) when illuminated with PFD 1920 μmol m^−2^ s^−1^ of blue/red, which is equal to approximately 417 W m^-2^ at leaf surface. Using photo-calorimetric measurements, we estimated that the power emitted from the illuminated leaves ranged from 103.3 to 111.0 W m^-2^ per leaf area (Table 1). Approximately from provided PFD of 1920 μmol m^−2^ s^−1^, of which about 1612 μmol m^−2^ s^−1^, 84% or ∼ 350 W m^-2^ was absorbed by the leaves. We calculated this with measurements of the PFD on the abaxial and adaxial side of a leaf and assumed a leaf reflection constant (ca. 3 %). From these, we calculated that the foliar heat emission of 103 -111 W m^-2^ (Table 1) was ca. 30 ± 7 % of the absorbed light, which was dissipated as heat via the NPQ and another quenching mechanism. These findings indicate that leaves emit a considerable amount of heat when exposed to moderate EAE.

**Table 1.** Heat emission from leaves of *Arabidopsis thaliana, Acer platanoides* and *Euphorbia pulcherrima* normalized *per* g of Fresh Weight (FW) or square meter of the leaf’s area. Measurements were made in a custom-made photo-calorimeter with constant photon flux density (PFD) of blue/red LED light of 1920 μmol m^-2^ s^-1^ or 417 W m^-2^ to submerged in water leaf. Mean values of calories and Joules are derived from 5 large leaves (*n*=5) or from 15 Arabidopsis rosettes (*n*=15).

| | $\text{Cal s}^{-1} \text{g}^{-1} (\text{FW})$ | $\text{J s}^{-1} \text{g}^{-1} (\text{FW})(\text{W g}^{-1})$ | $\text{Cal s}^{-1} \text{m}^{-2}$ | $\text{J s}^{-1} \text{m}^{-2} (\text{W m}^{-2})$ |
| --- | --- | --- | --- | --- |
| *Acer platanoides*<br>Sunnyside | $0,153 \pm 0,029$ | $0,643 \pm 0,125$ | $25,5 \pm 6,6$ | $106,7 \pm 27,9$ |
| *Acer platanoides*<br>Shade side | $0,148 \pm 0,039$ | $0,622 \pm 0,166$ | $24,6 \pm 7,9$ | $103,2 \pm 33,0$ |
| *Euphorbia pulcherrima* | $0,165 \pm 0,026$ | $0,691 \pm 0,111$ | $26,4 \pm 4,2$ | $110,6 \pm 17,8$ |
| *Arabidopsis thaliana* | $0,165 \pm 0,024$ | $0,693 \pm 0,100$ | $26,5 \pm 3,8$ | $111 \pm 16,1$ |

### Generic photoinhibition and NPQ reduction decrease foliar temperature

We then investigated whether NPQ and *F*_v_/*F*_m_ generic inhibition could lower leaf temperature under EAE stress. For the experimental setup, we treated Arabidopsis and maple leaves with the photosynthetic electron transport inhibitor 3-(3,4-dichlorophenyl)-1,1-dimethylurea (DCMU), which binds non-covalently to the plastoquinone (Q_B_) binding site at photosystem II, blocking electron transfer between Q_B_ and the plastoquinone pool. It is well established that this disruption of P680 charge separation strongly reduces NPQ (*8, 9, 22, 25*). We measured maximal foliar temperature differences (ΔT_max_) between non-saturating PFD (150 μmol m^−2^ s^−1^) and saturating PFD (4000 μmol m^−2^ s^−1^). In non-treated leaves, ΔT_max_ in maple and Arabidopsis was 23.1 ± 0.5 °C and 16.5 ±0.9 °C, respectively. Lanolin-coated leaves showed higher ΔT _max_ values (26.0 ± 0.7 °C and 18.6 ± 1.2 °C), consistent with impaired transpiration (Fig. 2), while DCMU-treated leaves exhibited a reduced ΔT_max_ (19.0 ±0.8 °C in maple and 12.4 ±0.8 °C in Arabidopsis, Fig. 3A and B). Moreover, the extent of ΔT_max_ suppression paralleled the DCMU concentration-dependent decreases in NPQ and *F*_v_/*F*_m_ (Fig. 3C and E). NPQ and *F*_v_/*F*_m_ were significantly reduced by DCMU but remained relatively unchanged in lanolin-treated leaves, indicating that photoinhibition of PSII differs mechanistically from transient transpiration inhibition in foliar thermoregulation. By changing DCMU doses, we established linear relationships between foliar temperature changes (ΔT_max_), reductions of NPQ (ΔNPQ) and *F*_v_/*F*_m_ (Δ*F*_v_/*F*_m_) in maple and Arabidopsis leaves (Fig. 3D and F). These linear correlations demonstrate that foliar heat production is proportional to the decline in NPQ and *F*_v_/*F*_m_ values.

**Figure 3.**
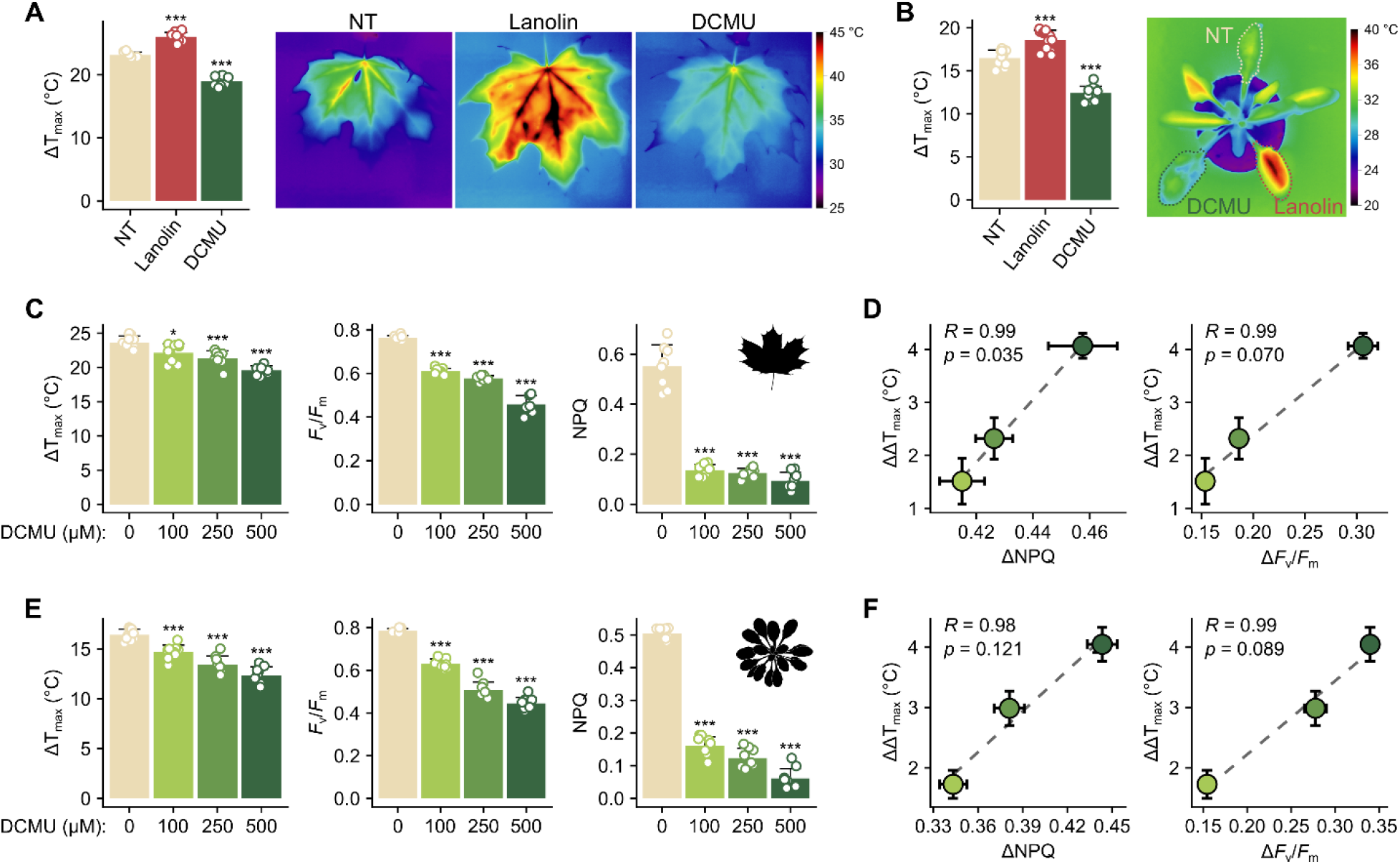
Effects of generic inhibition of the photosynthetic electron transport and inhibition of transpiration on foliar temperature. **(A, B)** Maximum foliar temperature increase (ΔT_max_) and representative thermographic images of untreated (NT), lanolin-coated, and (3-(3,4-dichlorophenyl)-1,1-dimethylurea) DCMU-treated leaves in *Acer platanoides* (A) and *Arabidopsis thaliana* (B). Data are mean ± SD (*n*=12) from three independent experiments. **(C, E)** Dose–response of ΔT_max_, maximal quantum yield of PSII (*F*_v_/*F*_m_), and non-photochemical quenching (NPQ) to increasing DCMU concentrations (0, 100, 250, 500 μM) in maple (**C**) and Arabidopsis **(E).** Data are mean ± SD (*n*=12). **(D, F)** Correlations between changes in ΔT_max_ (ΔΔT_max) and (i) ΔNPQ or (ii) Δ *F*_v_/*F*_m_ in maple **(D)** and Arabidopsis **(F)**; Pearson’s correlation coefficient (R) and p-value are indicated. Statistical significance in panels A and B (versus NT) and in panels C and E (versus 0 μM DCMU) was assessed by Tukey’s HSD test (* p < 0.05; *** p < 0.001).

To test whether generic partial inhibition of NPQ and *F*_v_/*F*_m_ can mitigate high light, thermal, and drought stress, we subjected Arabidopsis plants to eight days of drought in a control growth chamber. In the middle of the experimental setup, at day 4 a moderate EAE stress (600 μmol m^-2^ s^-1^) was imposed for the final four days in the presence or absence of DCMU (Fig. 4A). Under these conditions, DCMU treatment reduced foliar temperature and heat emission (Fig. 4C) by lowering NPQ and *F*_v_/*F*_m_ (Fig. 4D, E), without increasing plasma membrane damage, as shown by similar ion leakage in DCMU-treated and mock-treated plants (Fig. 4B). However, in contrast to DCMU-treated plants, control plants induced anthocyanins production, indicating for EAE stress in control plants but not in DCMU-treated plants (Fig S2).

**Figure 4.**
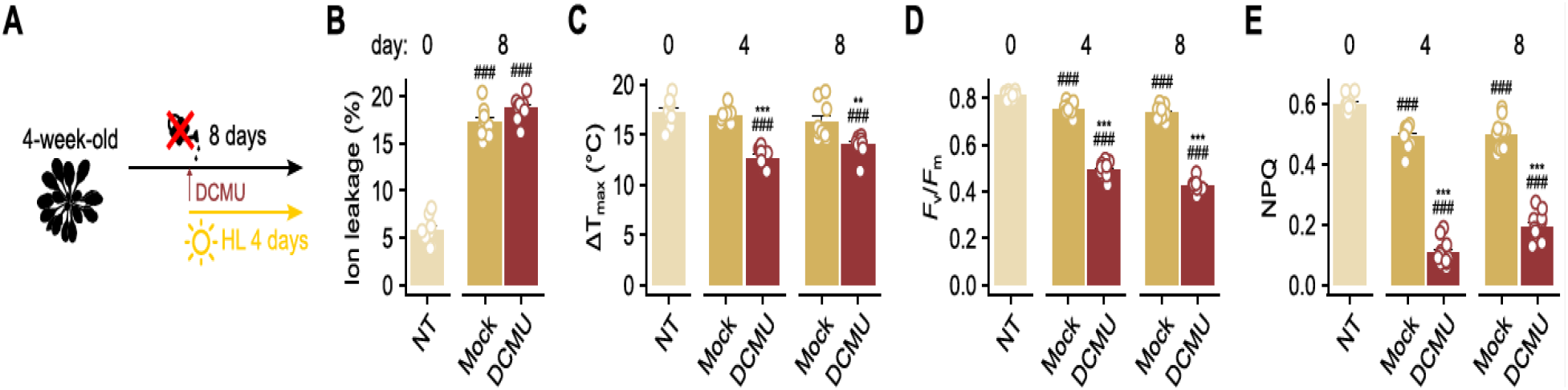
Effect of generic photoinhibition on foliar temperature and drought stress tolerance in Arabidopsis. Experimental timeline. Four-week-old *Arabidopsis thaliana* plants were subjected to an eight-day drought, with high light stress applied from day 4 to day 8, (3-(3,4-dichlorophenyl)-1,1-dimethylurea) DCMU or mock treatments were administered at the onset of high light (HL) stress with photon flux density (PFD = 600 μmol m^-2^ s^-1^) exposure applied to low-light adapted plants.(B) Relative ion leakage (%) measured on day 8 in untreated controls (NT), mock-treated, and DCMU-treated plants (mean ± SEM; *n*=9 from 3 biological repetitions). **(C–E)** Physiological responses in NT, mock-treated, and DCMU-treated plants on the 4^th^ and 8^th^ day (mean ± SEM; *n*=9, from 3 biological repetition): **(C)** Maximum foliar temperature increase (ΔT_max_), **(D)** Maximum quantum yield of PSII (*F*_v_/*F*_m_), and **(E)** Non-photochemical quenching (NPQ). Statistical significance was assessed by one-way ANOVA with Tukey’s HSD comparing NT (^#^ p < 0.05; ^##^ p < 0.01; ^###^ p < 0.001) and mock (* p < 0.05; ** p < 0.01; *** p < 0.001).

## Discussion

It is well known that plants generally cool down their surroundings through transpiration and reduce the atmospheric CO_2_ level due to photosynthesis (*17*). However, terrestrial plant leaves must lose water *via* transpiration to uptake atmospheric CO_2_ through stomata and must dissipate EAE as heat at the same time. The overall balance between photosynthesis, EAE dissipation, transpiration, and CO_2_ assimilation is complex and leaves conditionally, spatially and discreetly regulates stomatal conductance (transpiration) and CO_2_ assimilation (*30*). Therefore, a question arises about how leaves regulate their EAE dissipation as heat during variable EAE, photoinhibition, or transpiration inhibition.

Using thermal imaging (Fig. 1), we revealed a discrete, spatial, and emergent (wave-like), light- and age-dependent changes in foliar temperature (Figs. 2 and 3, Video S1 and S2) (*12, 13*). In Videos S1 and S2, it is shown that foliar temperature changes during constant EAE in Arabidopsis are indeed wave-like, which is consistent with previously observed changes in SAA and NAA (*9, 21–25, 30*). Under natural conditions, factors restricting transpiration, such as drought, EAE, and increased temperature, lead to enhanced NPQ and inhibit photosynthesis (Figs. 2 and 3) (*7, 31*). Our findings demonstrate that transpiration inhibition indeed leads to increased foliar temperature while partial inhibition of the photosynthetic electron transport and reduced NPQ shows the opposite effect (Figs. 2 and 3). Furthermore, a combined drought, high-light and DCMU treatment triggered inhibition of EAE-induced anthocyanin accumulation (Fig. 4, Fig. S2), thereby confirming of protection against the EAE stress. Taking into consideration the above data, which casts a new light on previous reports (*13, 18, 24, 25, 28, 29*), we concluded that reciprocated regulation of NPQ, transpiration, and photosynthetic electron transport controls not only plant productivity and stress tolerance but also foliar temperature. Thus, NPQ mechanism appears to have a dual role, both as a direct energy quenching and as a component of SAA and NAA signaling.

These experiments indicate that the differences in foliar temperature amplitude (ΔT_max_) between such conditions are up to 25% and demonstrate how conditionally foliar temperature, thus heat emission, can vary. We directly measured heat emission from leaves (Table 1, Fig. S1). All analyzed species released ca. 103.3–111.0 W m^−2^ as heat, approximately 30 ± 7% of the absorbed energy, substantially higher than previous theoretical estimates (*16*). Our results suggest that upper tree leaves (foliar surface carpet) directly exposed to sunlight under mild EAE stress in forest ecosystems could emit ca. 103–111 MW heat per km^2^. Taking into consideration our measurements of foliar heat emission and temperature amplitude in diverse conditions, total heat emission, for example, from 1,000,000 km^-2^ of upper leaves area in, for example, the Amazon rainforest can have conditional heat emission amplitude measured in TW scale. This can happen when transpiration and photosynthesis are reduced (7) due to high irradiation, high temperature, low relative humidity, and a reduction in soil water content in comparison to cloudy and high humidity days, and water available in the soil.

Although we cannot confirm whether such foliar conditional heat emission amplitudes operate at the global ecosystem scale, we can hypothesize, however, that variations in foliar heat emission driven by physiological changes in NPQ, photosynthetic electron transport, photorespiration, transpiration, and evaporation (*10, 14, 15, 20, 22, 23, 25, 26, 30*) may influence microclimates and conditionally contribute to the acceleration or inhibition of global warming. So far, this has not been investigated not only for the terrestrial plant ecosystems but also for the aquatic plant ecosystems. Therefore, the greenhouse effect models are inaccurate, and we are not able to fully understand why global warming is progressing.

## Acknowledgments

We are grateful to Prof. Frank Van Bresuegem from VIB and the University of Gent in Belgium for critical reading of the manuscript.

## Author contributions

SK proposed the working hypothesis and experimental design; RZG, MD, and MK performed the experiments; RZG analyzed data; RZG, MD, PB, PG, and SK wrote the manuscript; and SK supervised the research.

## Data and materials availability

All data are available in the main text or the supplementary materials.

## Methodology

### Plant Material and growing conditions

In this study, *Arabidopsis thaliana* wild-type (Col-0) and *Euphorbia pulcherrima* plants were cultivated in the growth chamber in a short photoperiod (9 h/15 h) under photon flux density (PFD) of 120 μmol photons m^−2^ s^−1^ at 22 °C and 70 ± 5% relative humidity. The mature green leaves of *Euphorbia pulcherrima* and 4-to 5-week-old *Arabidopsis* plants were used for analysis. The other plant species used in this study (*Acer platanoides, Quercus* sp., and hybrid Aspen (*Populus tremula x P. tremuloides*)) were grown outdoors in their natural habitat. The leaf samples from those plants were collected in May, June and October. For a combined drought/high-light stress experiment, Arabidopsis plants were not watered for 8 days. On day 4 half of the plants were sprayed with 500 μM DCMU, while the rest were mock-treated. Two hours after spraying all the plants dry out with Watman 3 MM and were exposed to PFD 600 μmol m^−2^ s^−1^ for 4 days in the same photoperiod.

### DCMU and lanolin treatment

The experiments described in this paper were performed as described before (involved two different treatments: chemical DCMU (3-(3,4-dichlorophenyl)-1,1-dimethylurea) treatment of leaves to block photosynthetic electron transport on the secondary electron acceptor Q_B_ and physical lanolin treatment to block stomatal conductance. For DCMU treatment, plants were sprayed with different concentrations: 100, 250, and 500 μM, while mock-treated plants were sprayed with water. After 3 hours, the plants were carefully dried with paper tissue and used in further experiments. During lanolin treatment, a thin layer of lanolin was gently applied to the abaxial side of the leaf to block stomata.

### Photo-calorimeter measurement

Calories produced by leaves were measured by a photo-calorimeter setup with two chambers, reference and measurement, covered with a light-reflecting surface (thermos-like) to optimize light distribution and minimize external temperature influence. Each chamber contained one liter of distilled water at 22°C. Several *Arabidopsis thaliana* rosettes or a single leaf of the other analyzed plants, were placed adaxially on a circular net and submerged 2-3 cm below the water surface in the measurement chamber. A magnet stirrer ensured a uniform mixing of water. The same conditions were used in the reference chamber, except that it contained no plant tissue. Both chambers were covered with thick plastic lids designed to hold thermo-sensors (Precision Measuring Instrument T995 instrument, Dostmann electronic GmbH, Wertheim, Germany), and a 1920 μmol m−2 s−1 blue/red LED light was provided to the submerged leaf surface. Light was evenly distributed in both chambers and reflected from the chamber glass-silver walls. Provided light intensity was measured on the adaxial and abaxial sides of leaves with a PFD light meter (SpectraPen SP, Photon Systems Instruments,Drasov, Czech Republic) with plastic lids covering chambers. Similarly, we measured PFD light on the abaxial and adaxial sides of tree growing leaves to get a clue how much energy is absorbed by leaves. Continuous temperature monitoring (around 30-40 minutes) was conducted *via* thermo-sensors connected to a data recording system connected to computer software, enabling the export of temperature changes every second for each chamber. The net temperature change generated by the sample (leaves) was determined by subtracting the reference chamber temperature changes from that of the measurement chamber. The heat emitted by the leaf was then calculated using water’s specific heat capacity and converted from calories to Joules to quantify the sample’s Watt output. Area and weight of leaves were measured and normalized *per* m^2^ or g FW.

### Foliar temperature measurements

The analysis was performed on whole soil-grown Arabidopsis rosettes using a FLIR A600-Series thermal imaging infrared camera. Determination of foliar temperature was performed using three different conditions:

- Variable light conditions: Measurements were based on an authorial program developed in the LabVIEW environment. Variable light conditions were applied as follows: 30 seconds of ambient light (150 μmol m^−2^ s^−1^), followed by 60 s of LED blue light (4000 μmol m^−2^ s^−1^), and then ambient light, which served as the recovery stage. The background temperature was subtracted from the maximum leaf temperature to calculate foliar temperature changes and eliminate the influence of environmental temperature. Analysis of thermograms was performed using OriginPro 8 software (FLIR Systems, Wilsonville, OR, USA). Thermographs of not detached tree leaves showing their foliar temperature on real time were taken using an FLIR T650sc IR camera with FLIR ResearchIR version 3.4 software (FLIR Systems, Wilsonville, OR, USA).
- Constant light conditions: the Arabidopsis rosette was exposed to 30 s of darkness, followed by 180 s of 2000 μmol m^−2^ s^−1^ white light, and again 30 s of darkness. Foliar temperature was analyzed using FLIR tools software.
- Increasing light intensity: a blue LED panel with various dark–light cycles: 120 s of darkness and 120 s of 500 μmol m^−2^ s^−1^ light, 60 s of darkness and 120 s of 1000 μmol m^−2^ s^−1^light, and 60 s of darkness and 120 s of 2000 μmol m^−2^ s^−1^ blue light. Foliar temperature changes of different leaf areas were measured and analyzed using FLIR Research IR version 3.4 software (FLIR Systems, Wilsonville, OR, USA).

### Chlorophyll *a* fluorescence

Chlorophyll *a* fluorescence parameters were determined on seedlings using a pulse amplitude-modulated FluorCam 800 MF and the associated software (Photon Systems Instruments, Drasov, Czech Republic). Before measurements, the plants were kept in darkness for a minimum of 30 min to determine *F*_0_ and *F*_m_, and then exposed to 5 min of actinic red light (90 μmol m^−2^ s^−1^) to determine *F*_t_ and *F*_m_’. The actinic light was switched off, and after incubation with far-red light, *F*_0_′ was determined. The effective photosystem II (PSII) (*F*_v_/*F*_m_) and NPQ were determined.

### Electrolyte leakage

The level of cell death in the DCMU-treated and high-light exposed plants was measured by electrolyte leakage. The whole rosette was removed and placed in 50 mL Falcon tubes containing 35 mL Milli-Q water. The relative electrolyte leakage was measured using a conductance meter (inoLab Cond Level 1, WTW, Weilheim, Germany) and calculated as a ratio of the value obtained after 1 hour incubation to the total leakage assessed after autoclaving the samples.

