## Supplementary file for "Transpiration, Photoinhibition and Non-photochemical Quenching Reciprocally Control Foliar Heat Emission"

**Supplementary Material:**

**Table S1**

**Figures S1-S2**

**Video S1 and S2:**

**Video S1 and S2:** Temporal profile of foliar temperature change (ΔT) in *Arabidopsis thaliana* leaves exposed to photon flux density (PFD) of 2000 μmol $m^{-2}$ $s^{-1}$. Videos were taken by two different FLIR cameras of two different experiments.

**Table S1**. $F_v/F_m$ and NPQ values measured in *Acer platanoides* leaves correspond to those shown in Figure 2, Panel C. Data are mean ± SD ($n$ = 6). Statistical significance was assessed by one-way ANOVA with Tukey's HSD comparing NT (* $p < 0.05$; ** $p < 0.01$; *** $p < 0.001$).

| Species | Treatment | Fv/Fm (mean ± SD) | NPQ (mean ± SD) |
|---|---|---|---|
| *Acer platanoides* (tree) | NT | 0.72 ± 0.053 | 0.60 ± 0.10 |
| | Lanolin (1h) | 0.70 ± 0.063 | 0.58 ± 0.140 |
| | Lanolin (2h) | 0.69 ± 0.054 * | 0.56 ± 0.13 * |

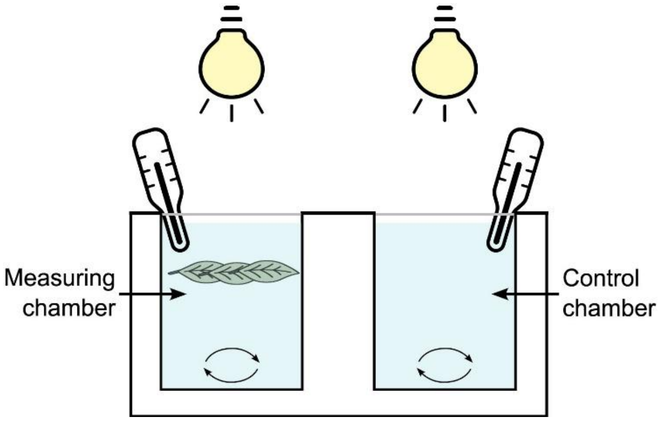


**Figure S1.** Schematic of the custom photo-calorimeter.

The instrument consists of two thermally insulated, stirred water baths: a measurement chamber containing the submerged leaf sample and a reference chamber without leaves. Each chamber is covered with a light-transmissive plastic cover to limit evaporation and equipped with a high-resolution temperature sensor. A controllable light source directs illumination, causing heat emission from the leaf surface. By continuously recording the temperature difference between the two chambers, the system achieves precise detection of subtle heat dissipation from the leaf.

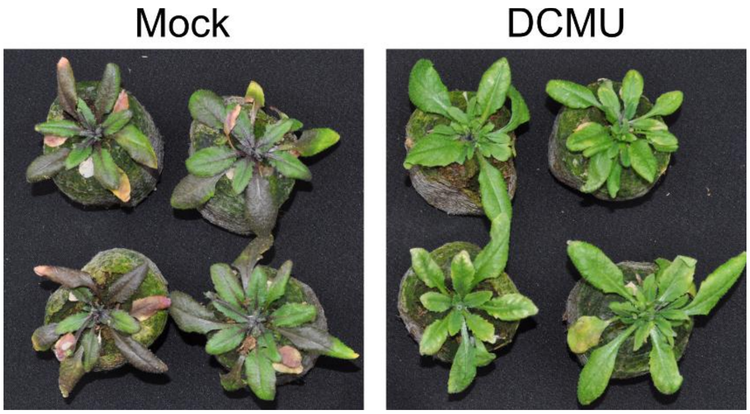


**Figure S2.** Phenotype of 3-(3,4-dichlorophenyl)-1,1-dimethylurea (DCMU) and mock-treated *Arabidopsis thaliana* plants exposed to combined drought and high light stress. Phenotypic characteristics of *Arabidopsis* rosettes after eight days of drought, with high light (600 µmol $m^{-2}$ $s^{-1}$) applied from day 4 to day 8. At day 4, plants were sprayed with 500 µM DCMU or mock-treated.
